# Drivers of the negative diversity-invasibility relationship: nutrient availablity, allelopathy, soil biota and soil legacy effects

**DOI:** 10.1101/2022.12.11.519939

**Authors:** Ayub M. O. Oduor, Mark van Kleunen, Yanjie Liu

**Affiliations:** Key Laboratory of Wetland Ecology and Environment, State Key Laboratory of Black Soils Conservation and Utilization, Northeast Institute of Geography and Agroecology, Chinese Academy of Sciences, Changchun, 130102, China; Department of Applied Biology, Technical University of Kenya, P.O Box 52428-00200, Nairobi, Kenya; Ecology, Department of Biology, University of Konstanz, 78464 Konstanz, Germany; Zhejiang Provincial Key Laboratory of Plant Evolutionary Ecology and Conservation, Taizhou University, Taizhou 318000, China

**Keywords:** biotic resistance, evolutionary novelties, exotic, global change, non-native, novel weapon, plant-soil feedback

## Abstract

Elton’s diversity-invasibility hypothesis predicts that high-diversity native communities should be less easily invaded than low-diversity communities. Although various mechanisms have been proposed to explain it, it remains unclear which of those mechanisms is more important and whether they operate simultaneously. Using one pool of native plant species and one pool of invasive alien plant species that naturally co-occur in China, we here tested in four separate experiments whether nutrient availability, allelopathy, soil microbiota and soil-legacy effects can all mediate the diversity-invasibility relationship. While soil-nutrient availability, allelopathy, soil biota and soil-legacy effects separately influenced biomass production of alien plant species and native plant communities, our results suggest that only soil biota and allelopathy influenced diversity-invasibility relationship in our study system. Importantly, by excluding the potential effects of allelopathy and soil biota in the nutrient-competition experiment, nutrient competition alone is not necessarily related to the negative diversity-invasibility relationship.

## INTRODUCTION

Invasions by alien plant species often reduce native biodiversity (Vila et al. 2011) and alter ecosystem processes (Mack et al. 2000). Successful establishment of an alien species in a site depends on various factors including the capacity of the species to invade (i.e., its invasiveness) and invasibility of the recipient native plant community (i.e., the resistance or susceptibility of the community to invasion) (Londsdale 1999). Elton’s diversity-invasibility hypothesis predicts that high-diversity communities should be less easily invaded than low-diversity communities (Elton 1958). Several empirical studies (mostly performed at small spatial scales such as plot levels and mesocosms) have found support for the diversity-invasibility hypothesis (Case 1990; Stohlgren et al. 1999; Fridley et al. 2007; Powell et al. 2011; Smith and Côté 2019; Zhang et al. 2020). Nevertheless, the mechanisms that underlie the diversity-invasibility relationship are not yet fully understood, although such knowledge is key to having a predictive understanding of invasibility of native plant communities.

Classical ecological theory predicts that resource-use complementarity is central to the resistance of high-diversity native communities to invasion (Case 1990; Byers and Noonburg 2003). Specifically, due to complementarity and selection effects, resident species in high-diversity communities may use limiting resources such as soil nutrients more completely and efficiently, leaving little available niche space for newly arriving species (Stachowicz et al. 1999; Loreau and Hector 2001; Kennedy et al. 2002). Moreover, high-diversity communities might also resist invasion because the communities may harbour competitively superior species that exclude invaders (Wardle 2001). The complementarity effect refers to resource partitioning or facilitative interactions between species, whereas the selection effect is caused by species that are particularly productive and competitive for resources in plant mixtures (Loreau and Hector 2001). Invaded terrestrial ecosystems often exhibit spatial heterogeneity in abiotic conditions including soil nutrients (Stohlgren et al. 1999; Fridley et al. 2007; Powell et al. 2011). Field observational studies have found that low-resource environments are generally less prone to invasion, with nutrient-rich habitats being the most invaded ones (Chytrý et al. 2008). Experiments have also shown that invasive plants often have greater growth and reproduction than native plants under high-nutrient conditions (Liu et al. 2017). Invasive plants benefit from an increase in resource availability because many of them have inherent fast growth strategies and the ability to rapidly exploit high-resource conditions that allow them to outcompete native plants (Dukes and Mooney 1999). Therefore, due to complementarity and selection effects, it is likely that negative diversity-invasibility relationship between native plant communities and invasive plants may occur more strongly in habitats with nutrient-poor soils than in habitats with nutrient-rich soils. Nevertheless, it remains unclear how variation in soil-nutrient levels (low vs. high) may influence resistance of native plant communities to plant invasion (Li et al. 2022).

Besides soil-nutrient availability, growth of plants in mixed-species systems is generally strongly influenced by soil allelochemicals (Inderjit et al. 2011; Zhang et al. 2021). Yet, the contribution of allelopathy to the resistance of species-rich native communities to invasion has received considerably less attention (Yuan et al. 2022). The homeland-security hypothesis predicts that native plants can resist invasion by producing allelochemicals that will deter growth of alien plant species (Cummings et al. 2012). Although only a few studies have tested this hypothesis, evidence is accumulating that some native species have allelopathic effects on alien species (Cummings et al. 2012; Christina et al. 2015; Mignoni et al. 2018; Yuan et al. 2021; Zhang et al. 2021). It is likely that species-rich native plant communities have stronger allelopathic effects on invaders than species-poor native communities (Yuan et al. 2022). This is because there may be a high likelihood that species-rich native communities harbour at least one species that produces a disproportionately potent allelochemical or there could be synergistic effects of different allelochemicals from different species within the community (Yuan et al. 2022). However, it remains unclear whether high-diversity native plant communities resist invasive plants more strongly through allelopathy than low-diversity native communities (Adomako et al. 2019; Yuan et al. 2022).

Diversity-invasibility relationship can also be influenced by soil biota (Zhang et al. 2020). Soil biota can strongly influence plant-community dynamics and may contribute to the coexistence of competing plant species or to the competitive dominance of one plant species over another (Batten et al. 2008). Soil biota influence plant growth and reproduction by acting as mutualists and antagonists of plants (Klironomos 2002; Callaway et al. 2004). Plant species diversity can differently influence activities and diversity of soil biota (Naylor et al. 2017; de Vries et al. 2019) that then have immediate consequences for plant growth. However, plants can also alter activities and diversities of soil biota that then feeds back to affect the plants’ growth and reproduction or that of newly arriving plant species – a phenomenon that is referred to as soil legacy effect (or plant-soil feedback) (Bever 2003). Soil legacy effect can be negative (a build-up of plant-specific pathogens), neutral (the net effect of the soil microbial community is zero), or positive (a plant accumulates mutualists that enhance its growth performance) (Bever 2003; Oduor et al. 2022). In line with the amplification-effect hypothesis, species-rich plant communities may accumulate a greater diversity and abundance of soil-borne pathogens than species-poor communities (Keesing et al. 2006), which then increases the odds that some of the pathogens will deter newly arriving species in the species-rich communities (i.e., negative soil-legacy effects of species-rich native plant communities) (Zhang et al. 2020). However, to date, few studies have explicitly tested the soil-legacy effects of native plant communities on alien plants (Müller et al. 2016; Heinen et al. 2018; Zhang et al. 2020).

Whether more than one mechanism that could underlie the diversity-invasibility relationship operate in the same invasion system has rarely been investigated (Mallon et al. 2015). Therefore, the goal of this study was to test whether competition for soil nutrients, allelopathy, soil biota and soil-legacy effects of native plant communities can all influence the diversity-invasibility relationship in the same study system. To test this, we used a study system of 14 native plant species and 20 alien plant species that co-occur in grasslands in China. Specifically, we performed four separate multispecies experiments to test the following predictions: 1) High-diversity native plant communities resist invasion by alien species more strongly than low-diversity communities in low-nutrient soils than in high-nutrient soils; (2) High-diversity native plant communities have stronger negative allelopathic effects on alien plant species than low-diversity native plant communities; (3) Soil biota influence the diversity-invasibility relationship; (4) Soils conditioned by high-diversity native plant communities have stronger suppressive effects on growth of alien plant species than soils conditioned by low-diversity communities.

## MATERIALS AND METHODS

### Study species and germination

To test the effects of soil nutrient availability, allelopathy, soil biota and soil legacy on the negative diversity-invasibility relationship, we conducted four multispecies experiments (**Fig. 1**). We used a total of 14 invasive alien and 20 native plant species (**Table S1**). We classified the species as invasive alien or native to China based on the book ―*The Checklist of the Alien Invasive Plants in China*‖ (Ma & Li, 2018) and the database Flora of China (www.efloras.org). To cover a wide taxonomic breadth, the 14 alien species were chosen from 12 genera of 7 families, while the 20 native species were chosen from 18 genera of 10 families. Seeds of the study species were either collected from wild populations or obtained from commercial seeds companies (**Table S1**). We raised seedlings from the seeds for use in the four separate experiments that were conducted in four compartments of a greenhouse at the Northeast Institute of Geography and Agricultural Ecology, Chinese Academy of Sciences (43°59’49”N, 125°24’03”E) as described below.

**Figure 1.**
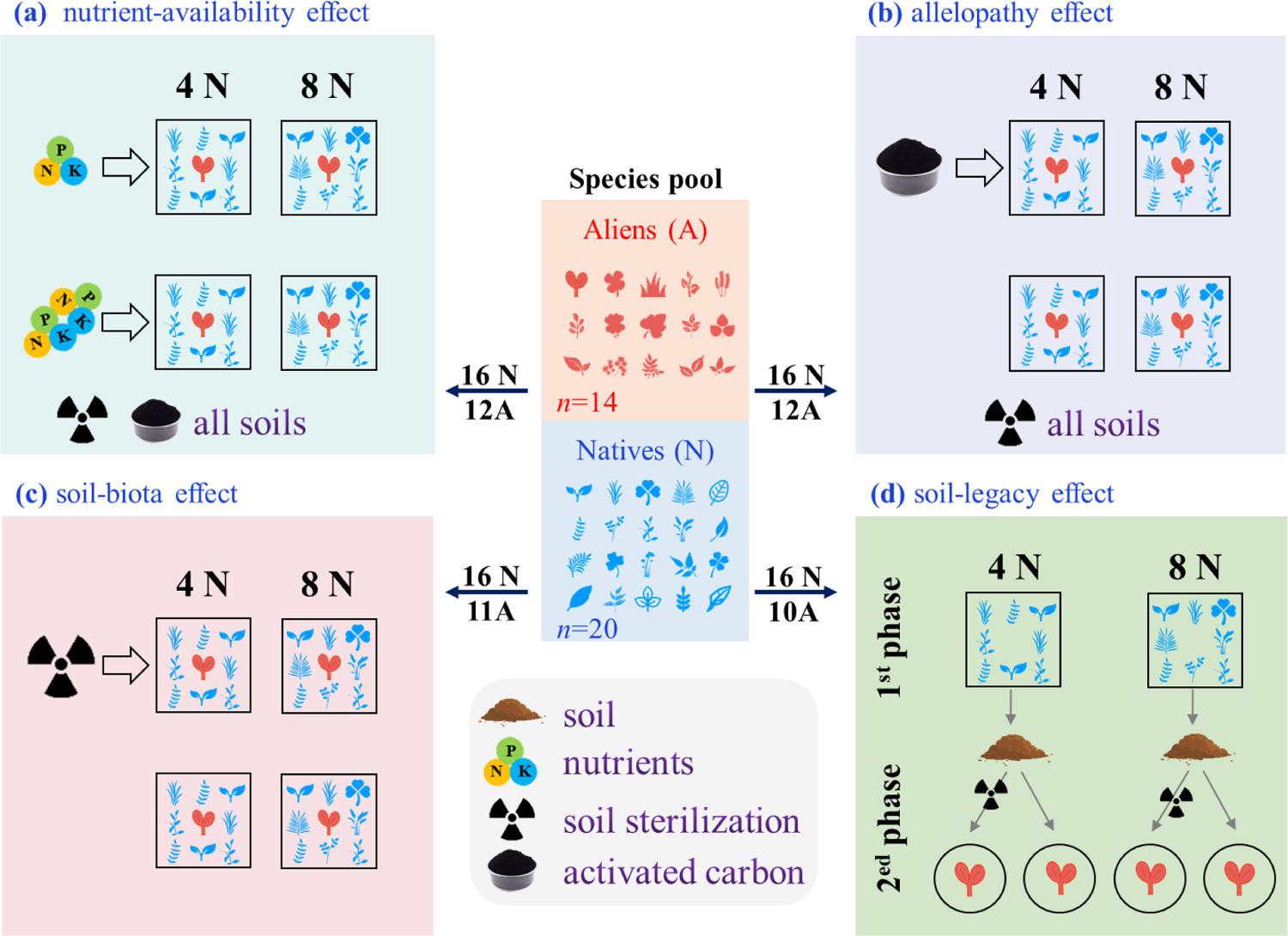
A schematic illustration of the experimental designs of the four separate experiments testing the nutrient-availability effect. (a), allelopathy effect (b), soil-biota effect (c) and soil-legacy effect (d) on the diversity-invasibility relationship.

Because the study species were known to have different germination rates (Yanjie Liu, personal observation), and the four experiments were set up on different days (9 April and 11 May 2021, respectively), we staggered sowing of seeds of the different species over various dates in each experiment (**Table S1**). We sowed seeds of the invasive alien and the native plant species into separate plastic trays (diameter × height: 25.5 cm × 4 cm) that had been filled with a sterilized peat-moss substrate (Pindstrup Plus, Pindstrup Mosebrug A/S, Ryomgård, Denmark). The substrate was sterilized with high pressure steam at 121 °C for 20 minutes to eliminate effects of soil biota. Due to the limited number of seeds for some species, the sets of study species used in the four experiments were not completely the same (**Table S1**; **Fig. 1**).

### Experiment 1: Do high-diversity native plant communities resist invasion more strongly than low-diversity native communities in low-nutrient soil?

#### Experimental setup

On 11 May 2021, we transplanted similar-sized seedlings of each species into 2.5 L circular plastic pots (top: 15.5 cm, bottom: 13.6 cm, height: 12.5 cm, 6 holes at the bottom) that had been filled with a 1:1 (v:v) mixture of washed sand and fine vermiculite. As we only aimed to test the effect of nutrient availability on the diversity-invasibility relationship in this experiment, we excluded the potential effects of soil biota and allelopathy. To exclude potential effects of soil biota, we sterilized the substrate with 25 kGy ^60^Coγ-ray irradiation at the CNNC Tongfu (Changchun) Radiation Technology Co., Ltd. in Jilin, China. To exclude potential allelopathic effects, we mixed 50 ml of activated carbon at a concentration of 20 ml / L with the sterilized substrate in all the pots.

We created eight low-diversity and eight high-diversity native plant communities in the individual pots from the pool of the 16 native species (**Table S2**). The low-diversity communities each had four native species, and the high-diversity communities each had eight native species (**Table S2**). Each species was used only twice for creating different low-diversity communities, and four times in the high-diversity communities (**Table S2**). For each of the 16 native communities, we introduced an individual seedling of one of the 12 alien species into the center of the pot (one alien species per pot). Within a pot, the plants were spaced at equal distances from each other. The plants in the different native community-alien plant combinations were then grown under two levels of nutrient treatment: low vs. high. In total, this experiment included 384 pots: 2 levels of native plant community diversity (low diversity vs. high diversity) × 8 community replicates × 2 nutrient levels (low-nutrient vs. high-nutrient) × 12 alien plant species (**Fig. 1a**). Immediately after transplant, we randomly assigned all the 384 pots to positions on four benches of a greenhouse, and all pots were re-randomized once during the experiment (i.e., 15 June 2021).

On 19 May 2021, eight days after transplant, we started weekly applications of nutrient treatments. For the low- and high-nutrient treatments, we applied 50 ml of a nutrient solution (Peters® Professional EC FERTILISER) at rates of 4 g/L and 1 g /L, respectively at weekly intervals for a total of seven weeks. The fertilizer contains N, P and K in the ratio of 20-20-20 and the following micro-nutrients: 0.02% Boron, 0.015% Copper, 0.12% Iron, 0.06% Manganese, 0.010% Molybdenum and 0.08% Zinc. The plants were watered regularly during the experiment. To avoid leakage of nutrient solution, we placed a plastic dish under each pot and re-applied the flowthroughs per pot to the respective pots.

On 10 July 2021, 60 days after transplant, we harvested all the experimental plants. For each pot, we separately harvested the above-ground biomass of the alien species and that of the native community. All above-ground biomass was dried to a constant weight at 65 ℃ for at least 72 hours and then weighed immediately to an accuracy of 0.001g. We then used the above-ground biomass to calculate biomass proportion of the alien species in each pot, applying the formula: above-ground biomass of alien species / [alien species + native community above-ground biomass]).

#### Statistical analysis

To test the effects of nutrient treatment, native community diversity, and their interaction on growth of alien plant species and native plant communities, we fitted linear mixed-effect models using maximum likelihood with the *lme* function in the *nlme* package (Pinheiro et al. 2007). In the models, above-ground biomass of the alien plant species, above-ground biomass of native communities, and proportional above-ground biomass of the alien species in each pot were treated as dependent variables. The native community diversity (low-diversity vs. high-diversity) and nutrient treatment (low-nutrient vs. high-nutrient) and two-way interaction between them were treated as fixed-effect independent variables. Species and family identities of the alien species and the community identity in which the alien species were grown were treated as random-effect independent variables. To assure normality of residuals and homogeneity of variance, above-ground biomass of the alien species and native community were natural-log-transformed, while the proportional biomass of the alien species in each pot was logit-transformed. We used the *varIdent* function to account for heteroscedasticity among the alien species for the models based on above-ground biomass and proportional biomass of the alien species. We also accounted for heteroscedasticity among the native communities for the model based on above-ground biomass of the native community. We tested the significance of the independent variables using likelihood-ratio tests (Zuur et al. 2009).

### Experiment 2: Do high-diversity native plant communities have stronger allelopathic effects on alien plant species?

#### Experimental setup

On 11 May 2021, we transplanted similar-sized seedlings of each species into 2.5 L circular plastic pots (same dimensions as used in experiment 1 above) that had been filled with a 1:1 mixture of sterilized washed sand and fine vermiculite. To exclude potential effects of soil biota and their interactions with allelochemicals, the substrate was sterilized with 25 kGy ^60^Coγ-ray irradiation at the CNNC Tongfu (Changchun) Radiation Technology Co., Ltd. in Jilin, China. In a half of the pots (n=192 pots), we homogenized the sterilized potting substrate with 50 ml of activated carbon to each pot (i.e. 20 ml activated carbon / L substrate). We created eight low- and eight high-diversity native plant communities (**Table S2**) as described above for experiment 1. We then planted one individual plant for each of the 12 alien species in the center of each pot (**Fig. 1b**). This resulted in 384 pots: 2 levels of native plant community diversity (low-diversity vs. high-diversity) × 8 community replicates × 2 levels of activated carbon presence in the pot (with vs. without) × 12 alien plant species.

Immediately after transplant, we randomly assigned all the 384 pots to positions on four benches of a greenhouse, and all pots were re-randomized once during the experiment (i.e., 15 June 2021). From 19 May 2021, eight days after transplant, we fertilized all the plants uniformly by adding 50 ml of a nutrient solution (Peters® Professional EC FERTILISER) at a concentration of 4 g/L into each pot at a weekly interval, for a total of seven applications. On 9 July 2021 (59 days after transplant), we first harvested and dried above-ground biomass of all the alien species and the native communities separately for each pot. We then calculated biomass proportion of the alien species in each pot as described above.

#### Statistical analysis

To test the effects of activated carbon, native community diversity, and their interaction on growth of alien plant species and the native communities, we fitted linear mixed-effect models using maximum likelihood with the *lme* function in the *nlme* package (Pinheiro et al. 2007). In the models, activated carbon, native plant community diversity, and the interaction between them were specified as fixed-effect independent variables, while above-ground biomass of the alien species, above-ground biomass of the native communities, and proportional biomass of the alien species in each pot were treated as dependent variables. To assure normality of residuals and homogeneity of variance, above-ground biomass of the alien species was natural-log-transformed, above-ground biomass of the native communities was square-root-transformed, and proportional biomass of the alien species in each pot was logit-transformed. To improve homoscedasticity of the residuals, we allowed the alien species and native community to have different variances by using the *varComb* and *varIdent* functions for the above-ground biomass of the alien species model. We also used the *varIdent* function to account for heteroscedasticity among the native communities for the above-ground biomass of the native community model. We tested the significance of the independent variables using likelihood-ratio tests (Zuur et al. 2009).

### Experiment 3: Do soil microorganisms influence the diversity-invasibility relationship?

#### Experimental set-up

To create live soil and sterilized soil treatments, we filled 2.5 L plastic pots (same volume as above) with a substrate mixture that comprised of 37.5% (v/v) sterilized sand, 37.5% (v/v) sterilized vermiculite and 25% (v/v) live or sterilized field soil. The live field soil served as an inoculum of a live soil microbiota and was collected from the top 20 cm layer at a grassland site in Jilin province in China (44°35′38”N, 125°30′54”E) on 2 April 2021. The field soil was sieved through a metal grid with a mesh size of 0.5 cm, and thereafter stored at 4 °C until use. Sterilization of the sand, vermiculite, and field soil was done with 25 kGy ^60^Coγ-ray irradiation at the CNNC Tongfu (Changchun) Radiation Technology Co., Ltd. in Jilin, China. Soil sterilization enabled us to compare the effects of live soil biota vs. non-live biota.

On 9 April 2021, we created eight low-diversity and eight high-diversity native plant communities in the individual pots from a pool of the 16 native species as described above (**Fig. 1c**; **Table S2**). We then transplanted individual similar-sized seedlings of the 12 alien plant species into the center of each pot. The experimental design resulted in 384 pots: 2 soil treatments (live soil vs. sterilized soil) × 2 levels of native community diversity (low diversity vs. high diversity) × 8 community replicates × 12 alien plant species). On 12 April 2021, we randomly assigned all the pots to positions on four benches of a greenhouse and re-randomized their positions once on 14 May 2021.

On 14 April 2021, five days after transplant, we started weekly additions of 50 ml of a nutrient solution (Peters® Professional EC FERTILISER) at a concentration of 4 g/L into each pot, for a total of seven rounds of application. On 4 June 2021 (56 days after transplant), we harvested all the experimental plants. For each pot, we separately harvested the above-ground biomass of the alien species and that of the native community. All above-ground biomass was dried to a constant biomass at 65 ℃ for 72 hours and then weighed immediately. We then calculated the proportional above-ground biomass of the alien species per pot as described above.

#### Statistical analysis

To test whether the presence of soil biota influenced the diversity-invasibility relationship, we fitted linear mixed-effect models using maximum likelihood with the *lme* function in the *nlme* package (Pinheiro et al. 2007). In the models, above-ground biomass of the alien plant species, above-ground biomass of the native plant communities, and proportional above-ground biomass of the alien species in each pot were treated as dependent variables. Native community diversity (low-diversity vs. high-diversity), soil treatment (sterilized soil vs. live soil) and two-way interaction between them were treated as fixed-effect independent variables. Species and family identity of the alien species and the community in which the alien species were grown were treated as random-effect independent variables. To assure normality of residuals and homogeneity of variance, above-ground biomass was cubic-root-transformed, while above-ground biomass of the native community and proportional biomass of the alien species were square-root-transformed. We used the *varIdent* function to account for heteroscedasticity among the alien species for all models. We tested the significance of the independent variables using likelihood-ratio tests (Zuur et al. 2009).

### Experiment 4: Do soils conditioned by high-diversity native plant communities have stronger suppressive effects on growth of alien plant species?

#### Soil-conditioning phase

On 2 April 2021, we collected field soil from the same site as above and sieved it through a 0.5 cm mesh. On 9 April 2021, we filled 2.5 L square plastic pots (same as above) with a substrate mixture of 37.5% (v/v) sand, 37.5% (v/v) vermiculite and 25% (v/v) live field soil (the same as in experiment 3). Prior to mixing with field soil, the sand and vermiculite had been sterilized (as in experiment 3). Then, we created eight low-diversity and eight high-diversity native plant communities in the individual pots from a pool of the 16 native species as described for the other experiments (**Fig. 1d**; **Table S1**). This design resulted in 144 pots: 2 levels of community diversity (low-diversity vs. high-diversity) × 8 community replicates × 9 replicates. On 12 April 2021, we randomly assigned all the pots to positions on two benches of a greenhouse. Position of the pots were re-randomized a month later (on 14 May 2021).

On 14 April 2021, five days after transplant, we started weekly additions of 50 ml of a nutrient solution (Peters® Professional EC FERTILISER) at a concentration of 4 g/L into each pot, for a total of 10 rounds of application. On 24 June 2021, 76 days after transplant, we harvested the plants. Immediately thereafter, we divided the soil in each pot into two portions: one portion was sterilized (as described above), while the other portion was left unsterilized (i.e., live soil) and stored at 4 ℃ in a refrigerator. The two soil portions were then used in the feedback-phase experiment described below. All above-ground biomass of the native communities was dried to a constant biomass at 65 °C for 72 hours and then weighed. The biomass was used as a co-variate in the statistical models described below.

#### Feedback phase

Between 23 May and 25 June 2021, we raised seedlings of nine alien plant species (**Table S1**) as described above. On 2 July 2021, we filled 144 pots with separate sterilized soils, and another 144 pots with live soils from the conditioning phase. The individual pots had a volume of 1 L (top diameter: 12.5 cm, bottom diameter: 9.5 cm, height: 11 cm). Immediately thereafter, we transplanted into the center of each pot one seedling of one of the 12 alien species (one individual per pot), which resulted in 288 pots: 2 levels of soil conditioning by the native plant communities (soil conditioned by low-diversity vs high-diversity) × 8 community replicates × 2 levels of soil sterilization treatment (sterilized vs live) × 9 alien plant species (**Fig. 1d**). Immediately after transplant, we randomly assigned all the pots to positions on four benches of a greenhouse, and all pots were re-randomized once during the experiment (i.e. on 27 July 2021).

On 6 July 2021, four days after transplant, we started weekly additions of 20 ml of a nutrient solution (Peters® Professional EC FERTILISER) at a concentration of 4 g/L into each pot, for a total of five rounds of applications. To avoid cross-contamination of the individual pots with soil microbes, we placed a plastic dish under each pot. On 13 August 2021, 42 days after transplant, we harvested all the experimental plants. For each pot, we separately harvested the above-ground and below-ground biomass. All the biomass was then dried to a constant weight at 65 °C for72 hours and then weighed. We calculated the total biomass (i.e., above-ground biomass + belowground biomass) and root mass fraction (i.e., above-ground biomass / total biomass) of each alien species per pot.

#### Statistical analysis

To test whether the alien species had lower growth in soils that had been conditioned by high-diversity native plant communities than by low-diversity native plant communities, we fitted linear mixed-effect models using maximum likelihood with the *lme* function in the *nlme* package (Pinheiro et al. 2007). In the models, total biomass and root mass fraction of the alien plant species were treated as dependent variables. The native community diversity (low-diversity vs. high-diversity) and soil type (sterilized vs. live soils that had been conditioned by the respective low and high-diversity native plant community communities) and the two-way interaction between them were treated as fixed-effect independent variables. Species and family identity of the alien species were treated as random-effect independent variables. We included biomass of the native plant communities as a co-variate to control for the potential differential effects of the communities on soil-nutrient content. To assure normality of residuals and homogeneity of variance, total biomass was natural-log-transformed, while root mass fraction was logit-transformed. We used the *varIdent* function to account for heteroscedasticity among the alien species for all models. We tested the significance of the independent variables using likelihood-ratio tests (Zuur et al. 2009).

## RESULTS

### Experiment 1: Do high-diversity native plant communities resist invasion more strongly than low-diversity native communities in low-nutrient soil?

Mean above-ground biomass of alien plant species (+244.7%) and native plant communities (+217.7%) were significantly higher in the high-nutrient than in the low-nutrient treatment regardless of the native plant community diversity levels (low diversity vs. high diversity) (**Table S3**; **Fig. 2a & b; Fig. S1a)**. A similar pattern was found for total above-ground biomass per pot (**Fig. S2a**). However, proportional above-ground biomass of the alien plant species was not significantly affected by nutrient treatment, native community diversity, and the interaction between them (**Table S3; Fig. 2c**).

**Figure 2.**
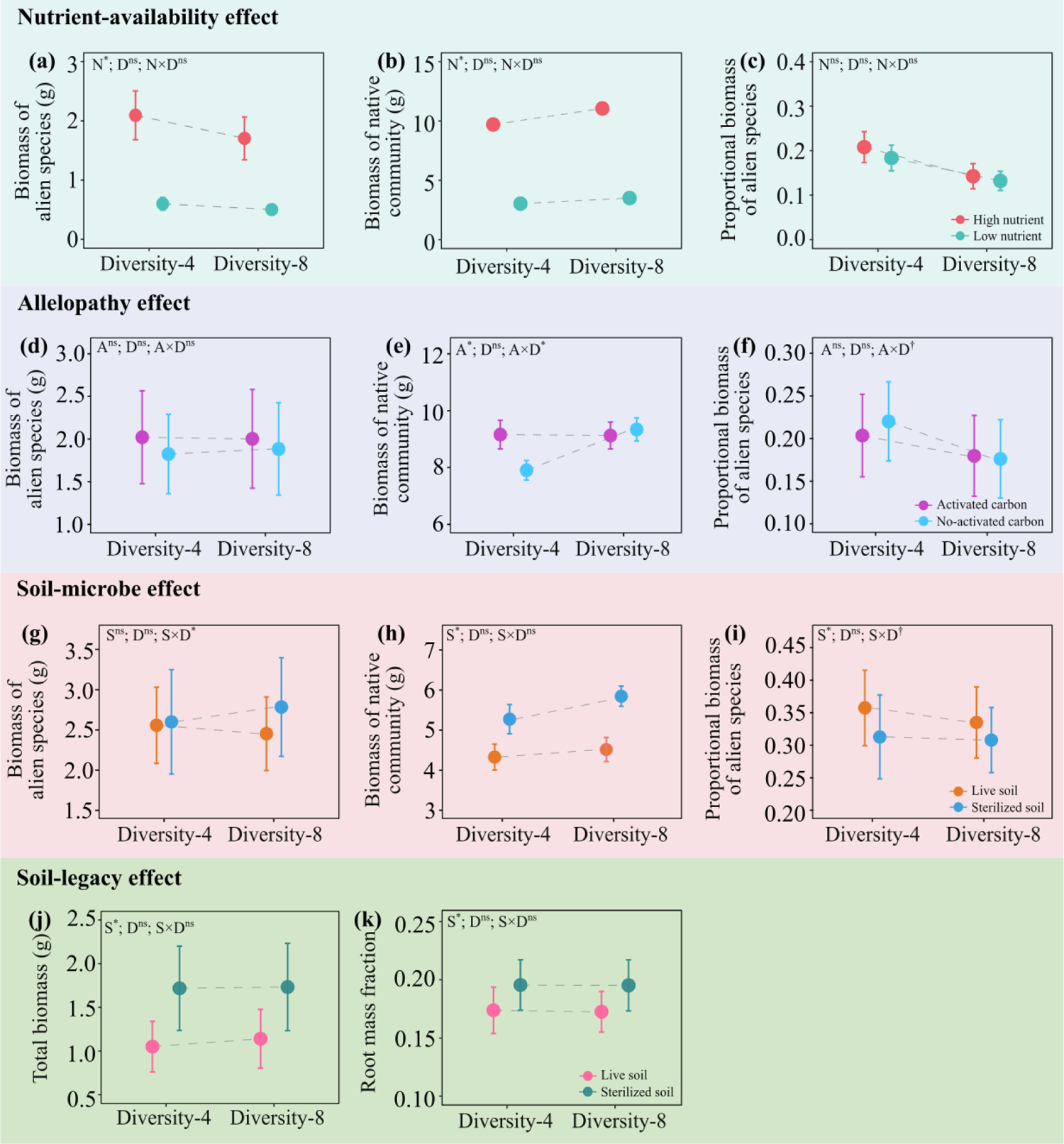
Mean (±1SE) values of total above-ground biomass of alien plant species. (**a, d, g**), total above-ground biomass of native plant communities (**b, e, h**), and proportional above-ground biomass of the alien species in each pot (**c, f, i)** under each treatment in separate experiments testing the nutrient-availability effect, allelopathy effect and soil-biota effect, as well as total biomass (j) and root mass fraction (k) of alien plant species in experiment testing the soil-legacy effect. Significant parameters (i.e., ‗N‘ [nutrient availability: low-nutrient vs. high-nutrient], ‗D‘ [diversity of native communities: diversity-4 vs. diversity-8],, ‗A‘ [activated carbon], ‗S‘ [soil sterilization], ‗N × D‘, ‗A × D‘ and ‗S × D‘) are indicted with asterisks (*: *P* < 0.05), marginally significant ones are marked with daggers (†: 0.05 ≤ *P* < 0.1), and the non-significant ones are marked with ‗ns‘.

### Experiment 2: Do high-diversity native plant communities have stronger allelopathic effects on alien plant species?

Mean above-ground biomass of the alien plant species was not affected by main and interactive effects of native community diversity and activated carbon treatment (**Table S4; Fig. 2d**). On the other hand, mean above-ground biomass of native communities was significantly influenced by the main effect of the activated carbon treatment (**Table S4; Fig. S1b**) and the interactive effect of native community diversity and activated carbon treatment (**Table S4; Fig. 2e**). Specifically, when grown without activated carbon, native communities produced more (+18.2%) above-ground biomass in high diversity than low-diversity conditions (**Fig. 2e**). However, community diversity level had minimal effects on above-ground biomass of the native communities when grown with activated carbon (a reduction in above-ground biomass by 0.4% in high-diversity communities relative to low-diversity communities) (**Fig. 2e**). A similar pattern was found for mean total above-ground biomass per pot (**Fig. S2b**). The mean proportional above-ground biomass of alien species per pot was marginally influenced by interactive effects of native community diversity and activated carbon treatment (**Table S4; Fig. 2f**). Specifically, high-diversity native communities tended to cause a stronger decline in proportional above-ground biomass of the alien species in the absence of activated carbon (−20.0%) than in the presence of activated carbon (−11.7%), relative to the effects of low-diversity communities (**Fig. 2f**).

### Experiment 3: Do soil microbiota influence the diversity-invasibility relationship between native plant communities and alien plant species?

Mean above-ground biomass of the alien plant species was significantly influenced by interactive effects of native plant community diversity and the soil-sterilization treatment (**Table S5; Fig. 2g**). Specifically, when grown in sterilized soil, the alien species increased their above-ground biomass yield (+7.1%) in high-diversity communities relative to when grown in low-diversity communities (**Fig. 2g**). However, an opposite pattern was observed in live soil as the alien species decreased mean above-ground biomass production (−4.1%) when grown in high-diversity communities relative to low-diversity communities (**Fig. 2g**). Soil sterilization significantly increased mean above-ground biomass of the native communities (+24.9%) and decreased mean proportional above-ground biomass of the alien species (−10.4%) (**Table S5; Fig. 2h; Fig. S1c**). Mean proportional above-ground biomass of the alien species was also influenced marginally by the interactive effect of native plant community diversity and soil sterilization (**Table S5**). Specifically, high-diversity native plant communities tended to cause a stronger decline in mean proportional above-ground biomass of alien plant species in live soil (−6.3%) than in sterilized soil (−1.5%), relative to the effects of low-diversity native communities (**Fig. 2i**). Mean total above-ground biomass per pot exhibited an opposite pattern to that of proportional above-ground biomass of alien species (**Fig. S2c**). In other words, the total above-ground biomass per pot tended to be higher for high-diversity communities than for low-diversity communities in sterilized soil (+8.7%) than in live soil (+1.2%).

### Experiment 4: Do soils conditioned by high-diversity native plant communities have stronger suppressive effects on growth of alien plant species?

Mean total biomass (−36.5%) and root mass fraction (−11.4%) of the alien species were significantly lower in live conditioned soil than in sterilized conditioned soil (**Table S6; Fig. 2j & k; Fig. S1d**). A similar pattern was found for above-ground and below-ground biomass (**Fig. S2d**). However, native community diversity level did not affect biomass of the alien species either separately or in interaction with soil type (**Table S6**). The native community biomass produced during the conditioning phase, which was included as a covariate in the models, did not have any significant effect on alien species biomass in the feedback phase (**Table S6**).

## DISCUSSION

Charles Elton proposed the diversity-invasibility hypothesis (Elton 1958) more than 60 years ago, and since then various mechanisms have been proposed to underlie the resistance of high-diversity native plant communities to invasion by alien plant. However, it remains unclear which of those mechanisms is more important and whether they operate simultaneously. Here, using one pool of native plant species and one pool of invasive alien plant species that naturally co-occur in China, we tested in four separate experiments whether nutrient availability, allelopathy, soil microbiota and soil-legacy effects can all mediate the diversity-invasibility relationship. Although nutrient availability, allelopathy, soil microbiota and soil-legacy effects separately influenced biomass production by alien plant species and native plant communities, our results suggest that only allelopathy (**Fig. 2**) and soil microbiota (**Fig. 2**) influenced the diversity-invasibility relationship in our study system.

### Experiment 1: Do high-diversity native plant communities resist invasion by alien species more strongly than low-diversity communities in low-nutrient soils?

Unexpectedly, we found that growth of the alien plant species was similar in the low- and high-diversity native plant communities regardless of the low- and high-nutrient treatment (**Fig. 2**). Based on Elton’s diversity-invasibility hypothesis (Elton 1958), we had predicted that high-diversity native plant communities would be more suppressive of alien plant growth than low-diversity native plant communities, and that this would be particularly the case at low nutrient availability. Our prediction was based on the idea that due to niche complementarity, diverse communities should better fill in the available niche space, and thus use more of the available resources (Loreau and Hector 2001). Niche complementarity can increase over time (Fargione et al. 2007; Jucker et al. 2020). Hence, it is possible that in the present study, resistance of the native communities against the alien invaders would have increased with diversity if our experiment would have lasted longer (the current experiment lasted 60 days). Indeed, long-term grassland biodiversity experiments have demonstrated that complementarity effects on productivity tend to strengthen through time (Cardinale et al. 2007; Zuppinger-Dingley et al. 2014; Guerrero-Ramírez et al. 2017). Therefore, long-term experimental studies are needed to test more rigorously the effect of soil nutrient availability on diversity-invasibility relationship and coexistence between alien plant species and native plant communities.

An alternative plausible explanation for the present results showing weak diversity-invasibility relationships is that the effect of the four-species native communities on the alien plant species was just as strong as that of eight-species native communities. A similar observation was previously made in two separate studies that varied species richness of native plants from 1, 2, 4, 6, 8, 12, to 24 species and found that a curve depicting performance responses of the invasive plants *Crepis tectorum* and *Digitaria ischaemum* to the various native species richness began to flatten out when the invaders were grown with six native species (Knops et al. 1999; Naeem et al. 2000).

More broadly, the present results complement those of other experiments that tested relationship between native plant community diversity and invasibility at small spatial scales (i.e., mesocosm and plot-level studies), with mixed findings. For example, separate experiments detected significantly stronger resistance of more diverse native plant communities to growth and reproductive performance of the alien plant invaders *Lolium temulentum* (Lyons and Schwartz 2001), *Cirsium arvense, Plantago major,* and *Agrostis stolonifera* (Levine 2001), *Crepis tectorum* (Naeem et al. 2000), *Lolium arundinaceum* (Rudgers et al. 2005) and *Centaurea maculosa* (Maron and Marler 2007). By contrast, other experiments did not find significant negative relationship between the diversity of the native plant communities and growth performance of alien plant species (Power and Sánchez Vilas 2020; Wei and van Kleunen 2022). The inconsistencies among the results that tested the diversity-invasibility hypothesis could be due to various factors including heterogeneity in resource conditions among the studies (Davies et al. 2005) and effects of microbial mutualists of plants in the invaded communities (Rudgers et al. 2005). Unlike previous studies, we excluded the potential effects of allelopathy (i.e., by mixing activated carbon in the substrate) and of soil biota (i.e., by sterilizing all substrate). Thus, the pattern we observed in the present study is likely driven only by competition for nutrients.

### Experiment 2: Do high-diversity native plant communities have stronger allelopathic effects on alien plant species?

We had expected that with increasing native plant-community diversity, the chance that some species within the communities would release strongly allelopathic compounds would increase or that allelochemicals of different species would act synergistically on the alien species. However, the finding that alien plant species had similar absolute above-ground biomass in low-diversity as in high-diversity native communities regardless of activated carbon treatment (**Fig. 2d**) suggests that the native plant communities did not exert significant allelopathic effects on the growth of alien species. A plausible explanation for the present finding would be that the alien species have developed adaptations to the allelochemicals produced by the native communities, due to the relatively long history (**Table S1**) of co-occurrence with the native plant species in China. Studies have shown that native plant species can adapt to competition from invasive alien plant species (Callaway et al. 2005; Oduor 2013, 2022) but the opposite is also likely to occur as some alien species may develop adaptations to native plant species. The present results corroborate those of Zhang et al. (2021), who conducted a meta-analysis based on 792 pairwise allelopathic effects of native plants on naturalized alien plants and found that such allelopathic effect is overall neutral (the net effect of the allelopathy is not significantly different from zero). Therefore, the idea that native plants can suppress alien plant growth by producing allelochemicals as proposed by the homeland-security hypothesis does not seem to find general support.

We found, however, that activated carbon addition tended to neutralize the positive effect of high diversity on above-ground biomass production of the native community (**Fig. 2e**). In other words, the native plant species likely had allelopathic effects on each other in low and not in high-diversity communities. This is also in line with the findings of Experiment-1 discussed above, in which we found that no matter the nutrient availability, diversity of native communities did not affect their above-ground biomass productions when growing in soils mixed with activated carbon. One plausible explanation for this finding is that in more diverse communities, the allelochemicals specific to each native species may be diluted to levels at which they are not yet effective (Yuan et al. 2022). Another likely explanation may be the reduction of self-inhibition through reduced release and increased degradation of autotoxic compounds in the high-diversity communities (*sensu* Xia et al. 2016). That is, if a species produces autotoxic allelochemicals when growing in monoculture, then presence of other species can cause a reduction in the release and an increased degradation of the autotoxic compounds (Xia et al. 2016). It is also possible that the alien species had allelopathic effects on the native plants, as found in the meta-analysis by Zhang et al. (2021), and that such negative allelopathic effects of the aliens on the natives were more likely compensated in highly diverse native plant communities.

Given that growth of the alien species was not influenced by allelopathic effects of native species, and only the native species had strong mutual allelopathic suppression on each other at low diversity, we consequently found tentative evidence that activated carbon addition ameliorated suppressive effects of high-diversity native communities on proportional above-ground biomass (i.e., dominance) of the alien species relative to those of low-diversity native communities. In other words, allelopathy plays, at least to some extent, a role driving the negative diversity-invasibility relationship by directly influencing the growth of native species rather than those of alien species.

### Experiment 3: Do soil biota mediate the diversity-invasibility relationship?

Live soil suppressed above-ground biomass of alien plant species in high-diversity native communities but not in low-diversity native communities (**Fig. 2g**), which supports our prediction that soil biota can influence the diversity-invasibility relationship. This prediction is further supported by our results showing that the high-diversity communities caused a marginally stronger decline in proportional above-ground biomass of the aliens in live soil than in sterilized soil (**Fig. 2i**). In other words, similar to allelopathic effects, soil biota also play a role in driving the negative diversity-invasibility relationship.

Plants can influence soil microbial communities through belowground carbon allocation, nutrient movement (Westover et al. 1997; Grayston et al. 1998; Dennis et al. 2010) and root exudates (de Vries et al. 2019) Consequently, plant-community composition can significantly influence the community composition of root-associated microbiota (Mitchell et al. 2010; Lange et al. 2015). It is generally accepted that soil microbial biomass, activity, and diversity can significantly increase with increasing plant diversity (Eisenhauer et al. 2010; Lange et al. 2015; Steinauer et al. 2015). The overall effect of soil-microbial communities (negative, neutral or positive) on plant growth may depend upon the balance of mutualists and pathogens present in a particular soil (van der Putten et al. 1993; Westover and Bever 2001; Klironomos 2002). In view of this, it is likely that the high-diversity native communities may have harbored a greater diversity and abundance of microbial antagonists of alien plants than the low-diversity communities (Hudson et al. 2006; Keesing et al. 2006; Zhang et al. 2020). Therefore, our finding indicates that the amplification effect (i.e., that native diversity is positively correlated with disease risk of aliens) (Keesing et al. 2006) may be a key driver behind the negative diversity-invasibility relationship.

In addition, our finding that live soil was generally more suppressive of native plant communities than sterilized soil (**Fig. 2**) suggests that the live grassland soil generally contained more microbial antagonists than mutualists of the native plants. Moreover, our tentative evidence that high plant diversity increased total above-ground biomass per pot more strongly in sterilized soils than in live soils (**Fig. S2c**) supports the idea that soil biota are important in mediating the diversity-productivity relationship in plant communities (Wang et al. 2019). However, our results also contradict a previous suggestion that species-rich plant communities should experience predominantly facilitative net effects by soil biota promoting plant growth, while species-poor plant communities are subject to antagonistic net soil effects due to the accumulation of pathogens (Eisenhauer et al. 2012). Clearly, further studies are still needed to explicitly test the absolute and relative roles of microbial antagonists and mutualists on the negative diversity-invasibility relationship, as well as on the positive diversity-productivity relationship.

### Experiment 4: Do soils conditioned by high-diversity native plant communities have stronger suppressive effects on growth of alien plant species?

Our finding that the alien plant species had similar growth in live soils that had been conditioned by low-vs. high-diversity native plant communities (**Fig. 2**) suggests that the two levels of native plant-community diversity had similar conditioning effects on the soil. These results contrast with those of other studies that have demonstrated significant effects of plant-community diversity on soil biological properties (Eisenhauer et al. 2010; Lange et al. 2015; Steinauer et al. 2015). Separate studies have shown that various factors can influence plant-diversity effects on soil microbial community functioning, including plant-species identity and richness, plant functional group richness, plant functional traits (Boeddinghaus et al. 2019; Sweeney et al. 2021), and duration of interactions between plants and soil (Kulmatiski and Beard 2008; Eisenhauer et al. 2010). For instance, various studies have reported a time-lag in plant-diversity effects on soil microorganisms and processes mediated by microorganisms (Hedlund et al. 2003; Bartelt-Ryser et al. 2005; Eisenhauer et al. 2010; Boeddinghaus et al. 2019). Thus, it is likely that we found similar soil-conditioning effects of low-diversity and high-diversity native plant communities partly because our soil condition phase of the experiment was conducted over a relatively short period of time – 76 days. Moreover, the significance of plant attributes in influencing soil-microbial communities may be context dependent and driven by edaphic (e.g., nutrient levels) and other environmental conditions (Eisenhauer et al. 2010; Tedersoo et al. 2016). Therefore, a difference in experimental conditions and study species may also explain the disparity between our finding and those of other studies that demonstrated significant correlations between plant species diversity and changes in soil functioning.

Our results complement those of the few other studies that investigated legacy effects of plant diversity on alien plant invasion, with mixed findings. For instance, soil legacy effects of an 8-species native plant community were similar to those of monocultures on the growth of an invasive plant *Bidens pilosa* (Xue et al. 2022). Another study, however, found that alien species experienced an 11.7% reduction in above-ground biomass when grown in soils trained by four native species than in soils trained by two native species (Zhang et al. 2020). The negative legacy effects in the latter study was attributed to a more diverse soil fungal and bacterial community in the soil conditioned by four species than in the soil conditioned by two species (Zhang et al. 2020). Plant-soil feedbacks have been studied extensively (van der Putten et al. 2013), but it remains unclear how soil-legacy effects of varying levels of native plant-community diversities affect plant community productivity in the wild due to a lack of studies (Heinen et al. 2020). In a recent study, plant communities with different combinations of grasses and forbs left soil legacies that negatively impacted succeeding plants of the same functional type (Heinen et al. 2020). More such studies that vary both native plant community functional diversity and species richness are required to provide more insights into soil-legacy effects of conditioning plant communities on productivity of responding plant communities.

## Conclusion

In conclusion, the present results suggest that in our study system of native grassland communities and alien plant species, soil biota and allelopathy may play a greater role than nutrient availability and soil-legacy effects in mediating the diversity-invasibility relationship. Importantly, by excluding the potential effects of allelopathy and soil biota in the nutrient-competition experiment, our results show for the first time that nutrient competition alone is not necessarily related to the negative diversity-invasibility relationship. We call for long term field-based experiments to help corroborate the present findings.

## Acknowledgements

We thank Huifei Jin, Yanjun Li, Mingxin Pan and Xue Zhang for help with the set-up of the experiment and plant harvest. We thank Yanmei Fu and Lichao Wang for help with visualization of the results. This work was supported by funding from the Chinese Academy of Sciences (Y9B7041001) and the Innovation Team Project of the Northeast Institute of Geography and Agroecology, Chinese Academy of Sciences (2022CXTD01). A.M.O.O acknowledges funding from the Chinese Academy of Sciences - President‘s International Fellowship Initiative (CAS-PIFI) (2021VBB0004).

## Author contributions

**Y. L** conceived the idea, designed and performed the experiment, and analyzed the data; **A.M.O.O** wrote the first draft of the manuscript, with further inputs from **Y. L** and **MvK**.

## Data accessibility

Should the manuscript be accepted, the data supporting the results will be archived in Figshare and the data DOI will be included at the end of the article.

## Supporting information

**Table S1.**
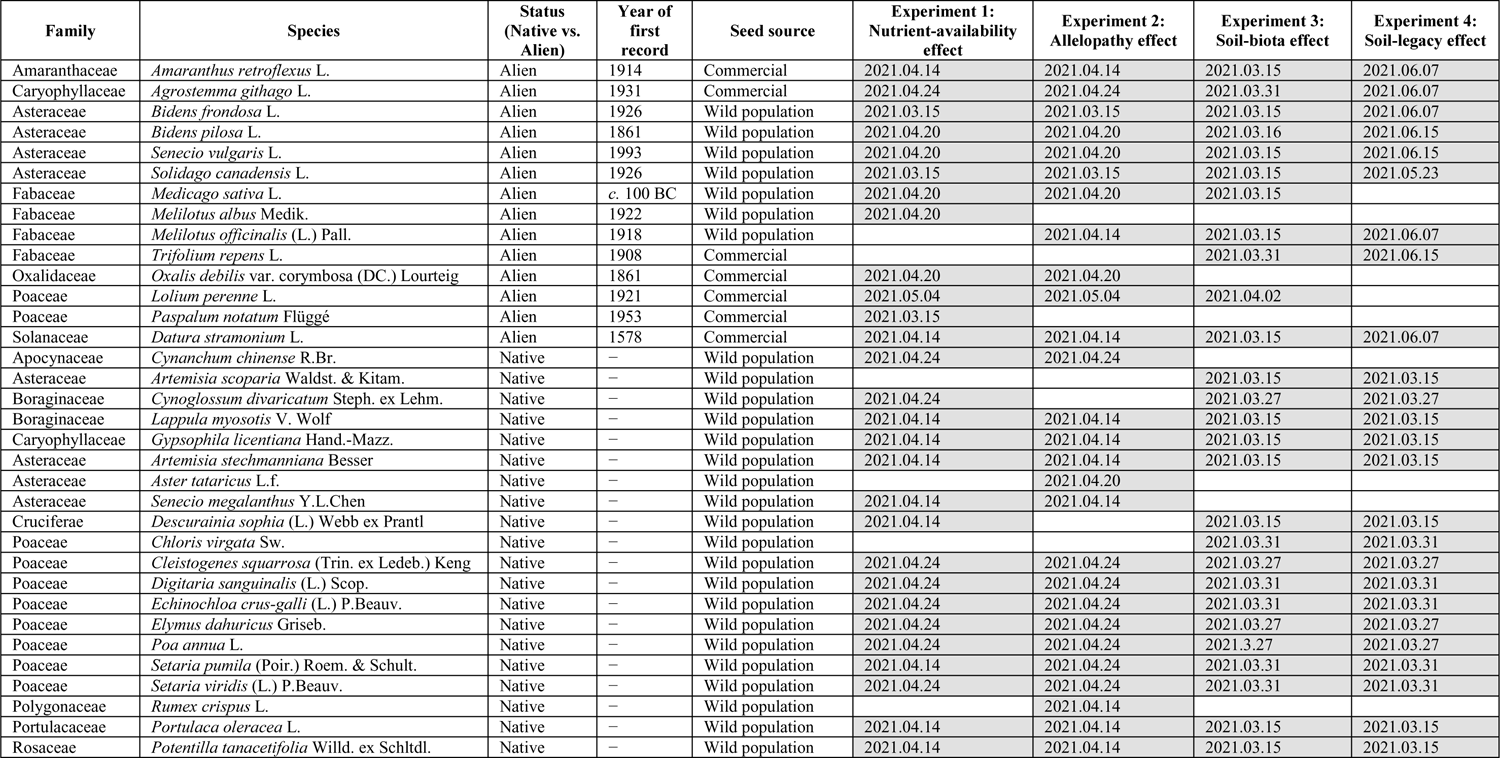
The 20 native and 14 alien plant species that were used in the four experiments (Experiment 1-Experiment 4). The species that were used in each experiment and their sowing dates are highlighted in grey.

**Table S2.**
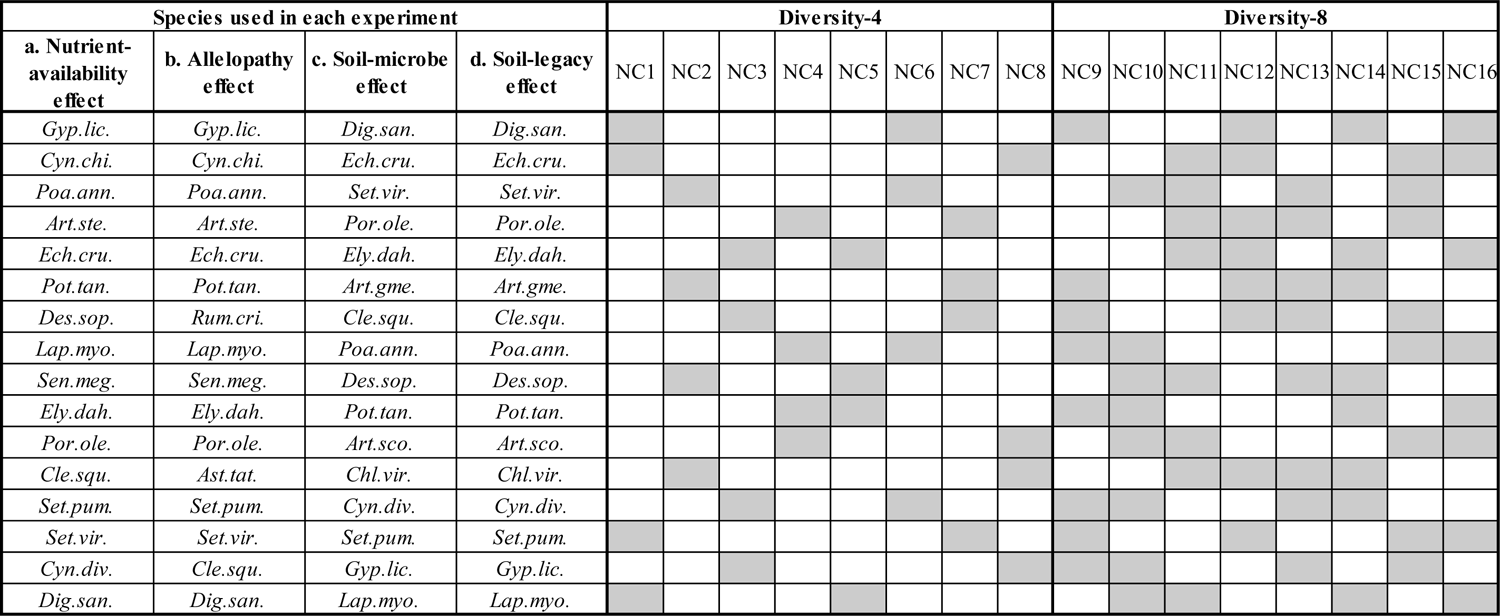
Compositions of 16 native plant communities (NC1-NC16), consisting of species that occur naturally in Chinese grasslands. The 16 communities were used in four separate experiments that tested effects of soil-nutrient availability, allelopathy, soil biota and soil legacies on the diversity-invasibility relationship. Eight communities (NC1 - NC8) had low diversity (four plant species), and the other eight (NC9 - NC16) had high diversity (eight plant species). Grey indicates the species present in the respective community. The names of the native species are abbreviated using the first three letters of the genus and the first three letters of the species epithet (for full species names, see Table S1).

**Table S3.**
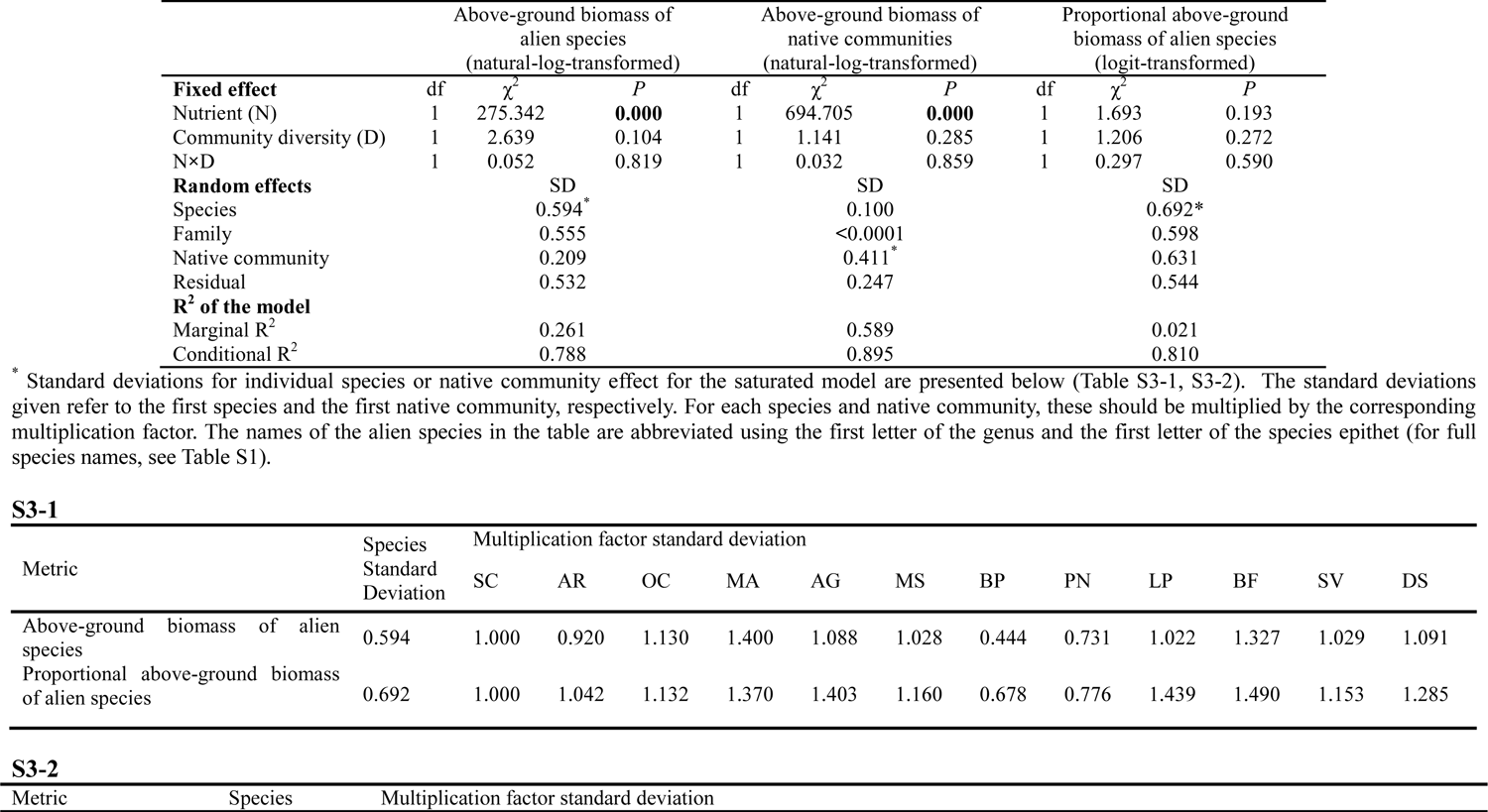
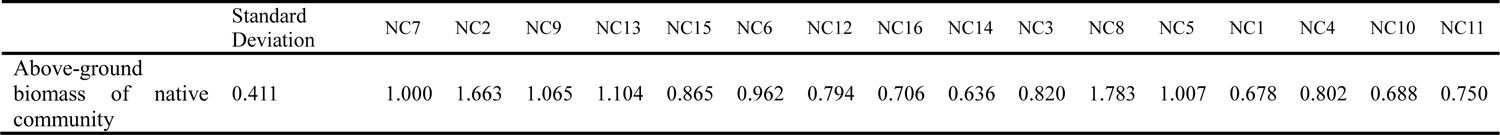
Results of linear mixed-effects models for experiment 1 that tested main and interactive effects of nutrient treatment (low-nutrient vs. high-nutrient) and native plant community diversity (low-diversity vs. high-diversity) on above-ground biomass of the alien plant species and native communities, as well as proportional above-ground biomass of alien plant species in each pot. Significant effects (*P* < 0.05) are in bold fonts.

**Table S4.**
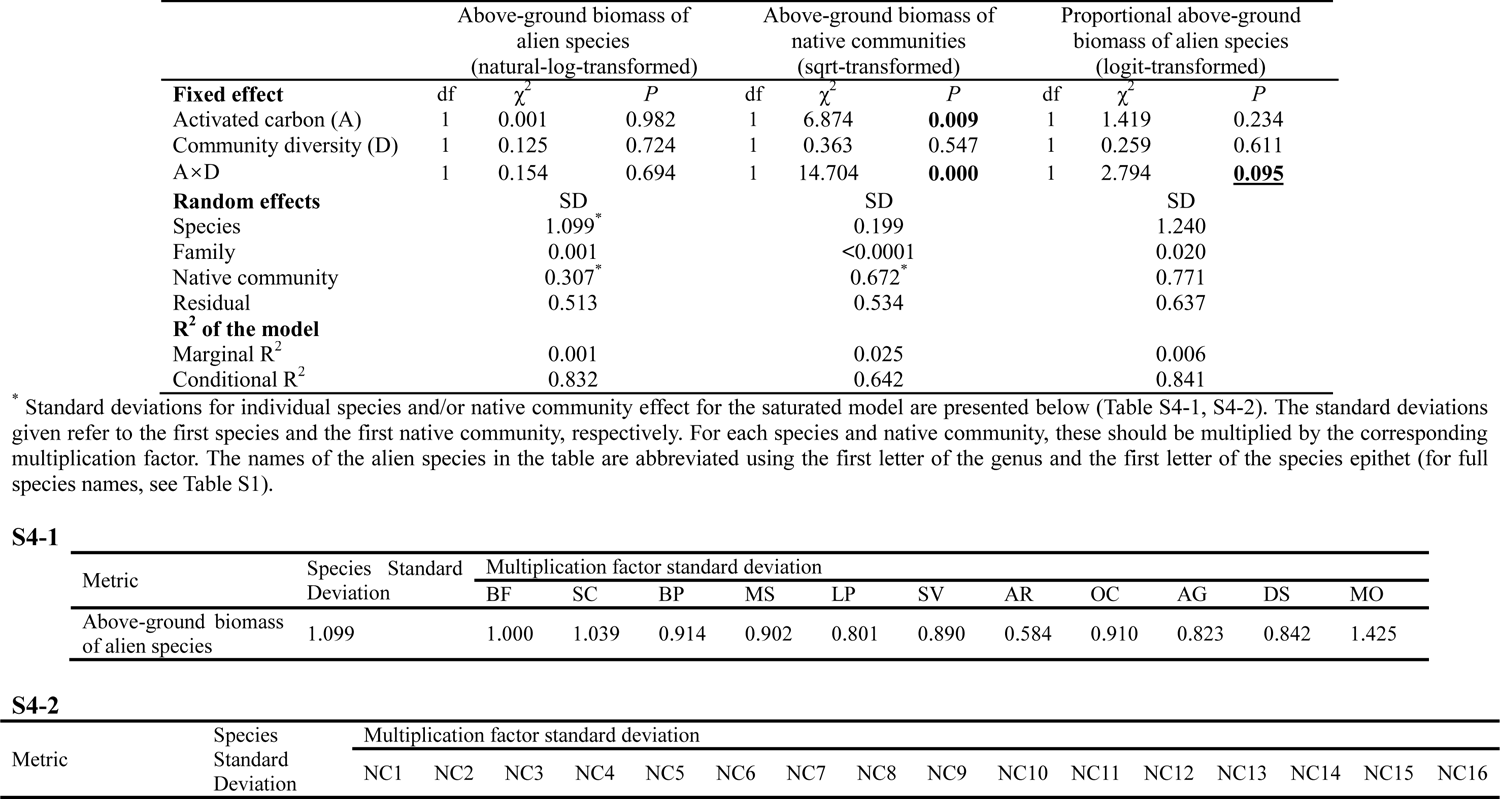
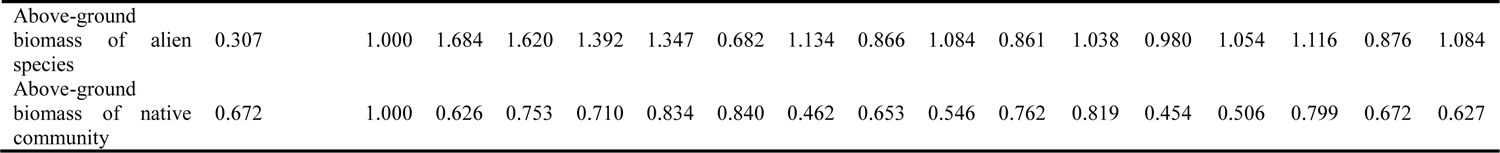
Results of linear mixed-effect models for experiment 2 that tested main and interactive effects of activated carbon (with vs. without activated carbon) and native plant community diversity level (low-diversity vs. high-diversity) on above-ground biomass of alien plant species and native communities, as well as proportional above-ground biomass of the alien plant species in each pot. Significant effects (*P* < 0.5) are highlighted in bold font, while marginally significant effects (0.05 < *P* < 0.1) are bolded and underlined.

**Table S5.**
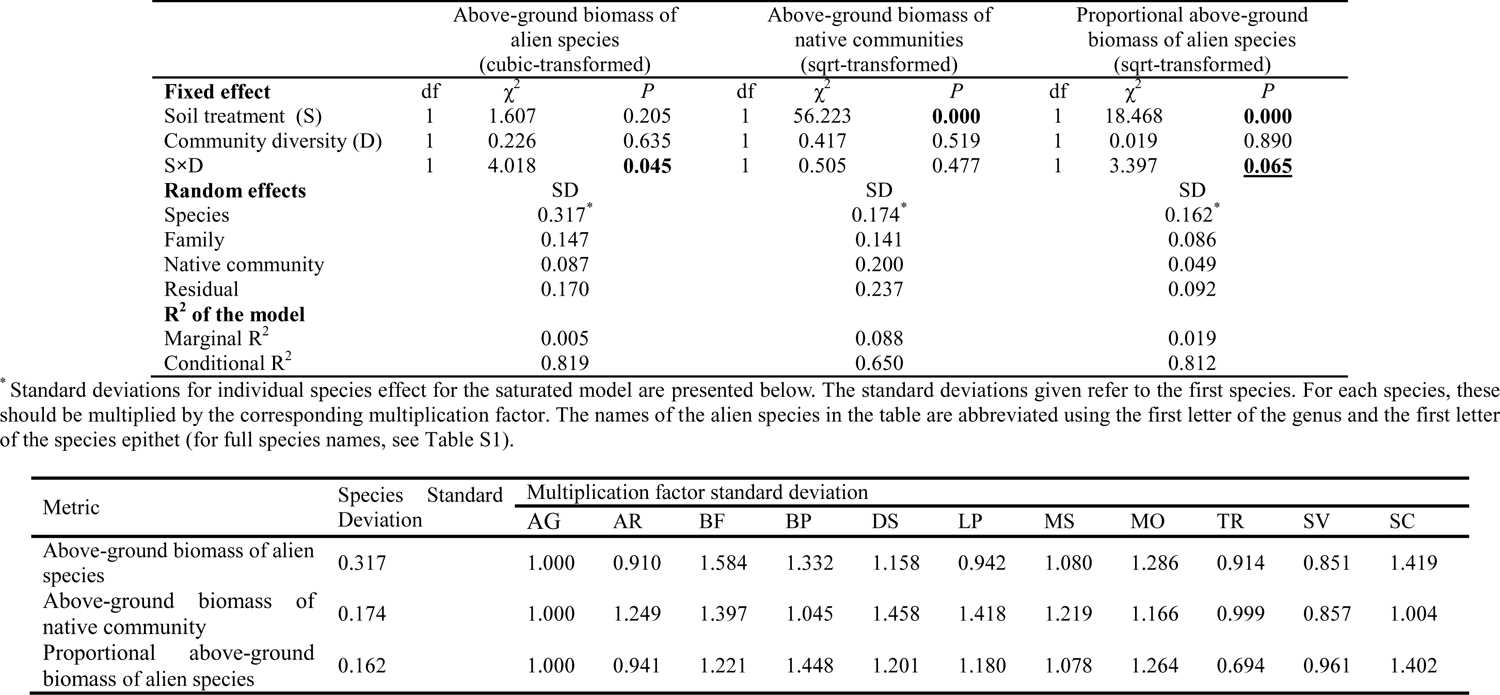
Results of linear mixed-effects models for experiment 3 testing the main and interactive effects of soil microorganisms (sterilized soil vs. live soil) and native plant community diversity (low diversity vs. high diversity) on above-ground biomass of the alien target plants and native community, as well as proportional biomass of the alien target species in each pot. Significant effects (P < 0.5) are highlighted in bold font, while marginally significant effects (0.05 < *P* < 0.1) are bold and underlined.

**Table S6.** Results of linear mixed-effects models for experiment 4 testing the main and interactive effects of soil-sterilization (sterilized soil vs. live soil) and native plant community diversity (low diversity vs. high diversity) of soil conditioning phrase on total biomass of the alien target plants and root mass fraction of the alien target species. Significant effects (P < 0.5) are highlighted in bold font.

|  | df | Total biomass<br>(natural-log transformed) |  | df | Root biomass fraction<br>(logit-transformed) |  |
| --- | --- | --- | --- | --- | --- | --- |
| | | $\chi^2$ | $P$ | | $\chi^2$ | $P$ |
| **Fixed effect** |  |  |  |  |  |  |
| Scale(log(Biomass of native community)) | 1 | 0.003 | 0.960 | 1 | 0.038 | 0.846 |
| Soil treatment (S) | 1 | 75.118 | **0.000** | 1 | 14.864 | **0.000** |
| Community diversity (D) | 1 | 0.064 | 0.800 | 1 | 0.075 | 0.785 |
| S×D | 1 | 1.765 | 0.184 | 1 | 0.006 | 0.939 |
| **Random effects** |  | SD |  |  | SD |  |
| Species |  | 0.590* |  |  | 0.439* |  |
| Family |  | 0.914 |  |  | 0.005 |  |
| Native community |  | <0.000 |  |  | 0.001 |  |
| Residual |  | 0.571 |  |  | 0.357 |  |
| **R<sup>2</sup> of the model** |  |  |  |  |  |  |
| Marginal R<sup>2</sup> |  | 0.036 |  |  | 0.013 |  |
| Conditional R<sup>2</sup> |  | 0.792 |  |  | 0.608 |  |

\* Standard deviations for individual species effect for the saturated model are presented below. The standard deviations given refer to the first species. For each species, these should be multiplied by the corresponding multiplication factor. The names of the alien species in the table are abbreviated using the first letter of the genus and the first letter of the species epithet (for full species names, see Table S1).

| Metric | Species Standard Deviation | Multiplication factor standard deviation |  |  |  |  |  |  |  |  |
| --- | --- | --- | --- | --- | --- | --- | --- | --- | --- | --- |
|  |  | AG | AR | BF | BP | DS | MO | TR | SV | SC |
| Total biomass | 0.590 | 1.000 | 0.333 | 0.983 | 0.774 | 0.889 | 0.853 | 0.948 | 0.893 | 0.915 |
| Root biomass fraction | 0.439 | 1.000 | 0.557 | 0.926 | 0.616 | 0.849 | 0.752 | 0.992 | 0.730 | 0.855 |

---

**Figure S1.**
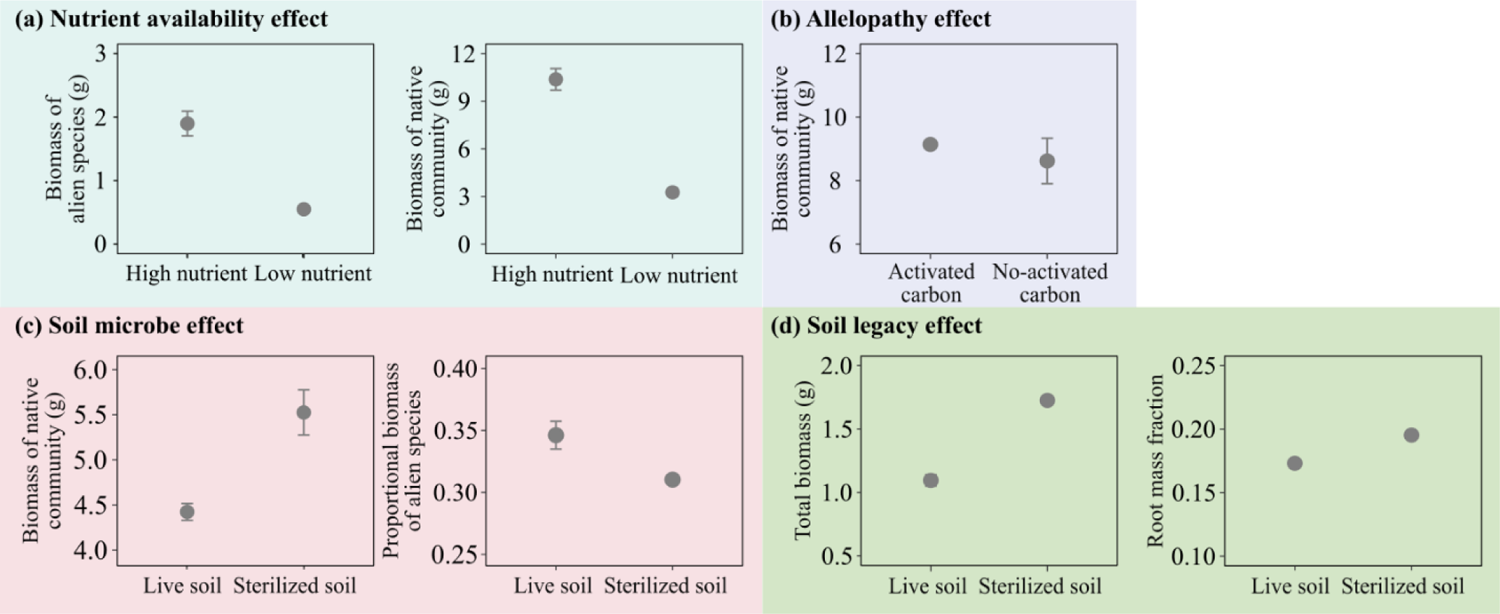
Mean (±1SE) values of above-ground biomass of alien plant species and native plant communities grown under low and high nutrient availability in experiment 1. (**a**), above-ground biomass of native communities grown with and without activated carbon in experiment 2 (**b**), above-ground biomass of native plant communities and proportional above-ground biomass of alien plant species in each pot in live and sterilized soil in experiment 3 (**c**), and total biomass and root mass fraction of the alien plant species grown in live vs. sterilized soils in experiment 4 (**d**).

**Figure S2.**
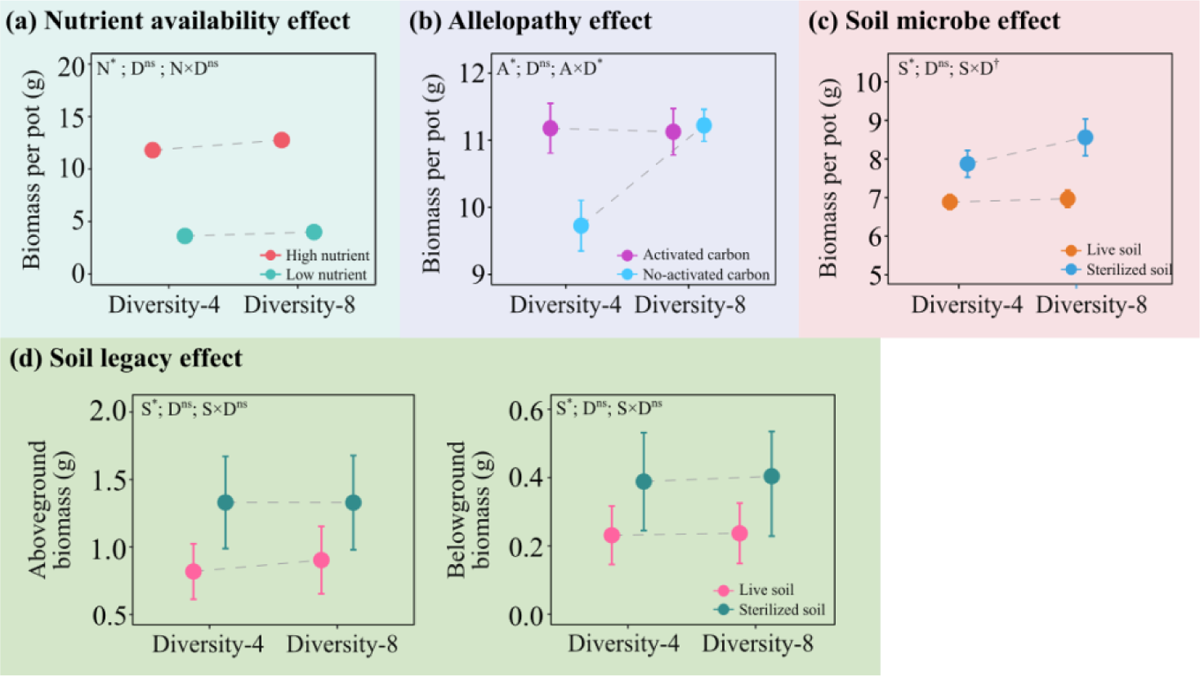
Mean values (±1 SE) of above-ground biomass per pot for experiment 1. (**a**), experiment 2 (**b**), and experiment 3 (**c**), as well as above- and below-ground biomass of alien plant species in experiment 4 (**d**). Significant parameters (i.e., ‗D‘ [diversity of native community], ‗N‘ [nutrient availability], ‗A‘ [activated carbon], ‗S‘ [soil sterilization], ‗N × D‘, ‗A × D‘ and ‗S × D‘) are marked with asterisks (*: *P* < 0.05), marginally significant effects are marked with daggers (†: 0.05 ≤ *P* < 0.1), and the non-significant ones are marked with ‗ns‘.

